# Widespread impact of DNA replication on mutational mechanisms in cancer

**DOI:** 10.1101/111302

**Authors:** Marketa Tomkova, Jakub Tomek, Skirmantas Kriaucionis, Benjamin Schuster-Böckler

**Author notes:** CORRESPONDING AUTHOR: Benjamin Schuster-Böckler Ludwig Cancer Research Oxford University of Oxford Old Road Campus Research Building Oxford OX3 7DQ United Kingdom.

## Abstract

DNA replication plays an important role in mutagenesis, yet little is known about how it interacts with other mutagenic processes. Here, we use somatic mutation signatures – each representing a mutagenic process – derived from 3056 patients spanning 19 cancer types to quantify the asymmetry of mutational signatures around replication origins and between early and late replicating regions. We observe that 22 out of 29 mutational signatures are significantly impacted by DNA replication. The distinct associations of different signatures with replication timing and direction around origins shed new light on several mutagenic processes, for example suggesting that oxidative damage to the nucleotide pool substantially contributes to the mutational landscape of esophageal adenocarcinoma. Together, our results indicate an involvement of DNA replication and associated damage repair in most mutagenic processes.

## Introduction

Understanding the mechanisms of mutagenesis in cancer is important for the prevention and treatment of the disease (Secrier et al. 2016; Stenzinger et al. 2014). Mounting evidence suggests replication itself contributes to cancer risk (Tomasetti & Vogelstein 2015). Copying of DNA is intrinsically asymmetrical, with leading and lagging strands being processed by distinct sets of enzymes (Lujan et al. 2016), and different genomic regions replicating at defined times during S phase (Fragkos et al. 2015). Previous analyses have focused either on the genome-wide distribution of mutation rate or on the strand specificity of individual base changes. These studies revealed that the average mutation frequency is increased in late-replicating regions (Stamatoyannopoulos et al. 2009; Lawrence et al. 2013), and that the asymmetric synthesis of DNA during replication leads to strand-specific frequencies of base changes (Shinbrot et al. 2014; Lujan et al. 2012; Reijns et al. 2015; Haradhvala et al. 2016). However, the extent to which DNA replication influences distinct mutational mechanisms, with their manifold possible causes, remains incompletely understood.

Mutational signatures have been established as a powerful approach to quantify the presence of distinct mutational mechanisms in cancer (Alexandrov et al. 2013). A mutational signature is a unique combination of the frequencies of all base-pair mutation types (C:G>A:T, T:A>G:C etc) and their flanking nucleotides. Since it is usually not known which base in a pair was the source of a mutation, the convention is to annotate mutations from the pyrimidine (C>A, T>A, etc.), leading to 96 possible combinations of mutation types and neighboring bases. Non-negative matrix factorization is used to extract mutational signatures from somatic mutations in cancer samples (Alexandrov et al. 2013). This approach has the important advantage of being able to distinguish between processes that have the same major mutation type (such as C>T transitions), but differ in their sequence context. We built upon this feature of mutational signatures and developed a computational framework to identify the replication-strand-specific impact of distinct mutational processes. Using this system, we quantified the replication strand and timing bias of mutational signatures across 19 cancer types. We show that replication affects the distribution of nearly all mutational signatures across the genome, including those that represent chemical mutagens. The unique strand-asymmetry and replication timing profile of different signatures reveal novel aspects of the underlying mechanism. For example, we discovered a strong lagging strand bias of T>G mutations in esophageal adenocarcinoma, suggesting an involvement of oxidative damage to the nucleotide pool in the etiology of the disease. Together, our results highlight the critical role of DNA replication and the associated repair in the accumulation of somatic mutations.

## Results and Discussion

### Replication bias of mutational signatures

DNA replication in eukaryotic cells is initiated around replication origins (ORI), from where it proceeds in both directions, synthesizing the leading strand continuously and the lagging strand discontinuously (Fig. 1A). We used two independent data sets to describe replication direction relative to the reference sequence, one derived from high-resolution replication timing data (Haradhvala et al. 2016) and the other from direct detection of ORIs by short nascent strands sequencing (SNS-seq) (Besnard et al. 2012), corrected for technical artefacts (Foulk et al. 2015) (see Methods). The former provides information for more genomic loci, while the latter is of higher resolution. As a third measure of DNA replication, we compared regions replicating early during S phase to regions replicating late (Haradhvala et al. 2016). We calculated *strand-specific* signatures (Morganella et al. 2016) that add strand information to each mutation type, based on the direction of DNA replication (Haradhvala et al. 2016) (Fig. 1B). We further condensed the strand-specific signatures into *directional signatures* consisting of 96 mutation types, each assigned either “leading” or “lagging” direction depending on the frequency in the strand-specific signature (Fig. 1C). These directional signatures can be used to separately compute the presence to the signature on the leading and lagging strand in individual samples, which we call the *exposure* to the signature in a sample (Fig. 1D). Depending on whether the strand bias matches the consensus of the directional signature, the exposure can be *matching* or *inverse.* The latter can occur if the strand bias of a signature in a subset of samples does not match the bias observed in the samples that most strongly contributed to the definition of the signature. We applied this novel algorithm to somatic mutations detected in whole-genome sequencing of 3056 tumor samples from 19 cancer types (Supplementary Table 1). We excluded protein-coding genes from the analysis in order to prevent potential confounding of the results by transcription strand asymmetry (Haradhvala et al. 2016; Alexandrov et al. 2013) or selection. Samples with microsatellite-instability (MSI) and POLE mutations were treated as separate groups, since they are associated with specific mutational processes. In total, we detected 25 mutational signatures that each corresponded to one of the COSMIC signatures^1^ and 4 novel signatures, which were primarily found in samples that had not been previously used for signature extraction (myeloid blood, skin, MSI, and ovarian cancers) (Fig. 1–figure supplement 1–6).

**Fig. 1:**
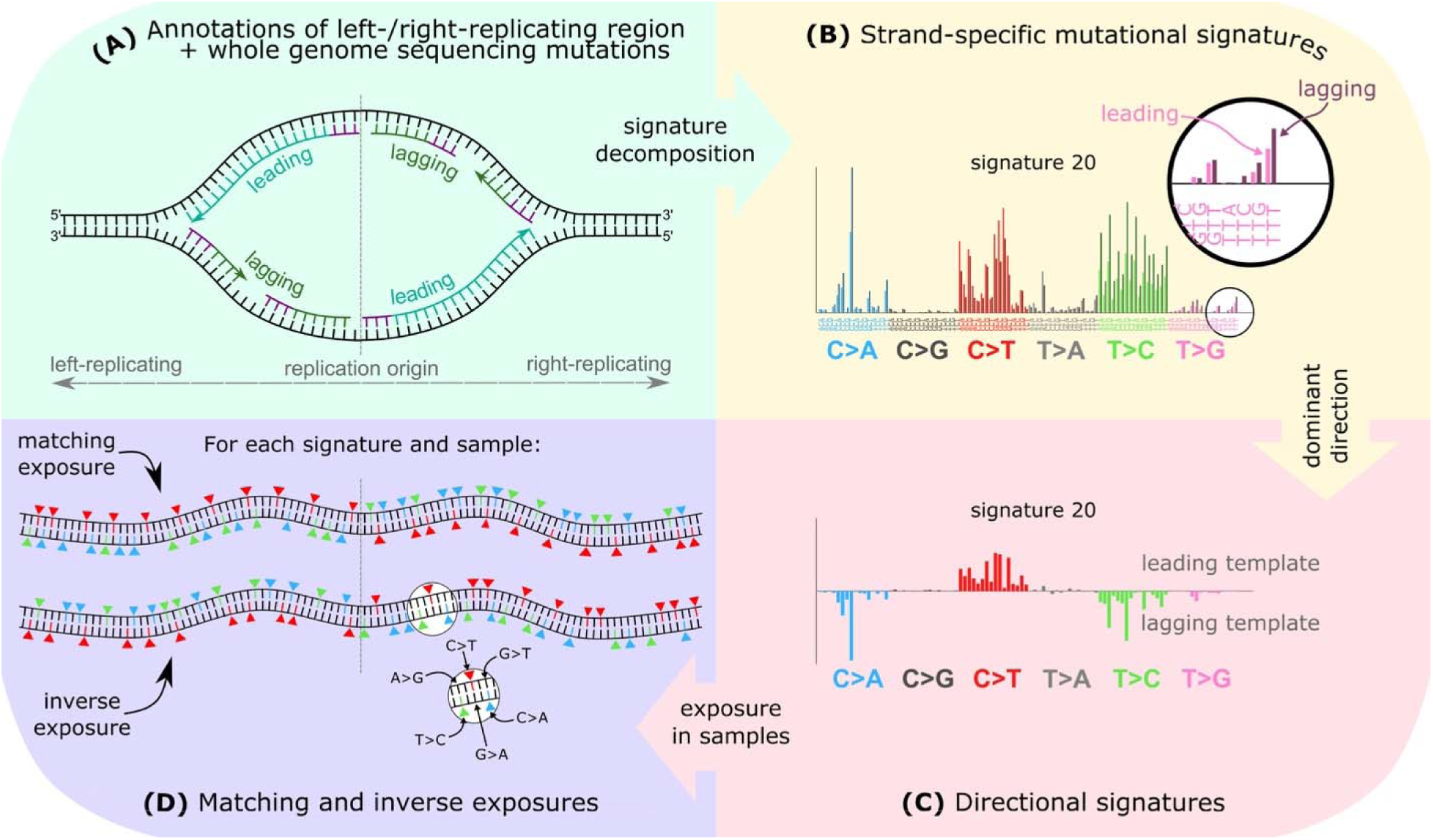
Methods overview. (A) Mutation frequency on the leading and lagging strand is computed using annotated left/right-replicating regions and somatic single-nucleotide mutations oriented according to the strand of the pyrimidine in the base-pair. **(B)** Leading and lagging strand-specific mutational signatures are extracted (signature 20 is shown as an example). **(C)** Each of the 96 mutation types is annotated according to its dominant direction (upwards-facing bars for leading, downwards-facing bars for lagging template preference). **(D)** Exposures to the directional signatures are separately quantified for the leading and lagging strand of each patient. The exposure in the *matching orientation* reflects the extent to which mutations in pyrimidines on the leading (and lagging) strand can be explained by the leading (and lagging) component of the signature, respectively. Conversely, the exposure in the *inverse orientation* reflects how mutations in pyrimidines on the leading strand can be explained by the lagging component of the signature (or vice-versa) (Methods). Top part of 1D shows an example of a sample with completely matching exposure, given the signature in 1C, with C>T mutations on the leading template and C>A and T>C mutations on the lagging template, whereas bottom part of 1D shows an example of a sample with completely inverse exposure.

In total, 22 out of 29 signatures exhibited significant replication strand asymmetry or significant correlation with replication timing (signtest p < 0.05, with Bonferroni correction; Fig. 2, Fig. 2–figure supplement 1, 4–figure supplement 1–10). Such widespread replication bias across the mutational landscape is surprising, considering that previous reports documented strand bias for only a few mutational processes such as activity of the APOBEC class of enzymes that selectively edit exposed single-stranded cytosines on the lagging strand (Morganella et al. 2016; Hoopes et al. 2016; Haradhvala et al. 2016; Green et al. 2016; Seplyarskiy et al. 2016). Including protein coding genes did not qualitatively change the results (Fig. 2–figure supplement 2, 5), nor did the exclusion of non-coding in addition to protein-coding genes (Fig. 2–figure supplement 3, 6). Similarly, using SNS-seq data to determine replication strand direction leads to highly similar findings (Fig. 2–figure supplement 4, 7). Furthermore, while the timing of replication of certain genomic regions is highly tissue-specific, we do observe a very strong correlation between replication timing data derived from different cell types, suggesting that tissue-differences will not fundamentally alter the results of this analysis.

**Fig. 2:**
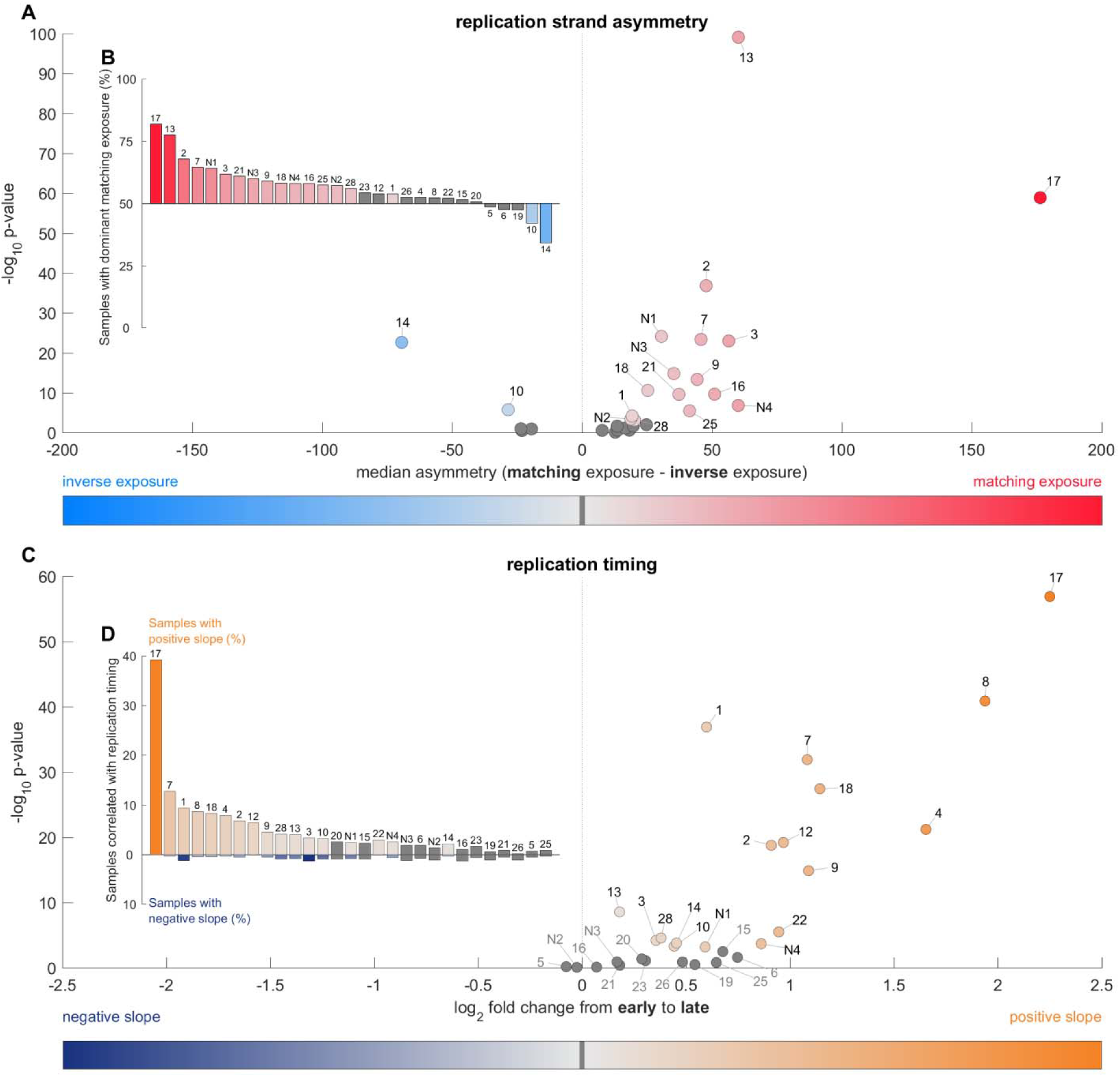
Most mutational signatures exhibit a significant replication strand asymmetry and/or correlation with replication timing. (A) The difference of matching and inverse exposure is computed for each sample and signature. For each signature, the median value of these differences (in samples exposed to this signature) is plotted against-log_10_ p-value (signtest of strand asymmetry per sample; with Bonferroni correction). **(B)** Percentage of samples that have higher matching than inverse exposure to the signature denoted above/below each bar. **(C)** Correlation of exposures with replication timing. The 20 kbp replication domains were divided into four quartiles by their average replication timing (early-replicated in the first quartile, late-replicated in the last quartile) and exposures to signatures were computed in each quartile. Log_2_-transformed fold change from average exposure in early (first quartile) to late (last quartile) is plotted on the x-axis, i.e. values on the right denote more mutations in late-replicated regions, values on the left reflect more mutations in early replicating regions. The y-axis represents significance of the direction of the correlation of signature with replication timing in individual samples (signtest of correlation sign per sample: 0 for non-significant correlation,-**1** for negative correlation, **1** for positive correlation; with Bonferroni correction). **(D)** Percentage of samples with a significantly positive and negative correlation with exposure, respectively.

Our observations confirm that both APOBEC signatures (2 and 13) exhibit clear strand asymmetry, with signature 13 being the most significantly asymmetric signature (p < 8e^-100^). We also observe differences in these signatures with respect to replication timing: signature 2 shows clear enrichment in late replicating regions (log_2_ fold-change 0.91 from early to late), whereas signature 13 shows only a mild increase in late replicating regions (log_2_ fold-change 0.18; Fig. 3), which is consistent with previous reports (Morganella et al. 2016). These results validate that our approach is able to correctly identify strand and timing asymmetries of mutagenic processes. Consequently, we next tried to interpret the replication biases we observed in other mutational signatures.

**Fig. 3:**
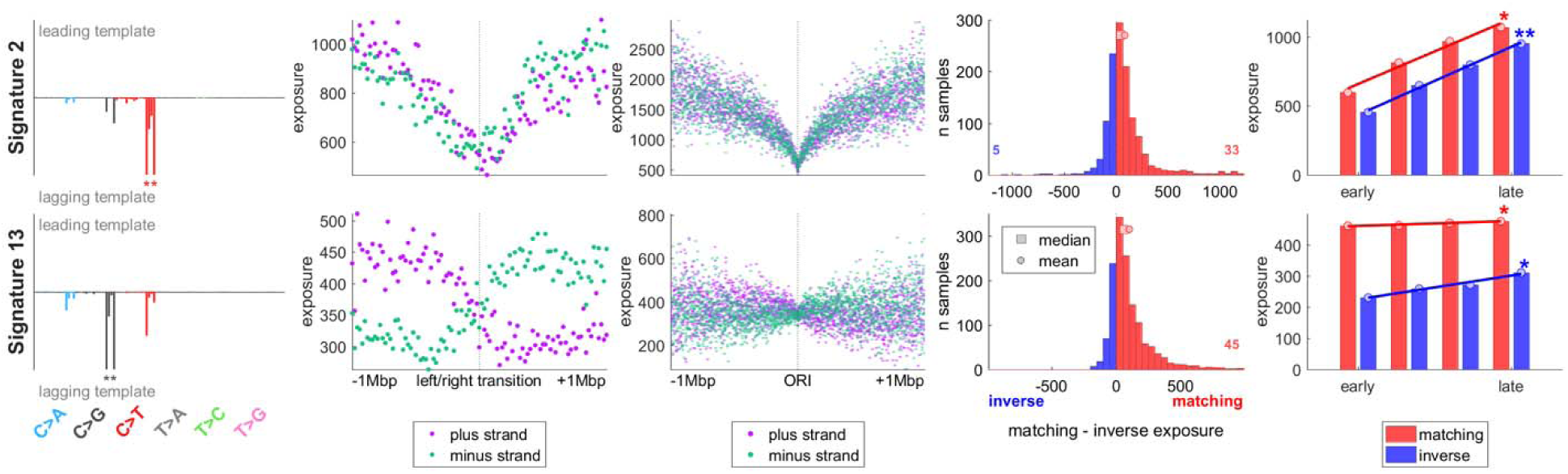
APOBEC signatures show strong but distinct effects of replication. Column 1: directional signatures for the two APOBEC signatures. Column 2: mean exposure on the plus (Watson) and minus (Crick) strand around transitions between left- and right-replicating regions. The transition corresponds to a region enriched for replication origins. Column 3: mean exposure on the plus and minus strand around directly ascertained replication origins. Column 4: distribution of differences between matching and inverse exposure amongst patients with sufficient exposure. Number of outliers is denoted by the small numbers on the sides. Column 5: mean matching and inverse exposure in four quartiles of replication timing; asterisks represent significance of the fit (F-test for coefficient of deviation from 0; ***P < 0.001; **P < 0.01; *P < 0.05). The leading and lagging strand annotations used in columns 4 and 5 are based on the direction of replication derived from replication timing data.

### Processes directly involving DNA replication or repair

Amongst the better understood mutational mechanisms, several involve replicative processes and DNA repair, such as mismatch-repair deficiency (MMR) (Zhao et al. 2014) or mutations in the proofreading domain of Pol ε (“POLE-M samples”) (Shinbrot et al. 2014; Shlien et al. 2015). We first analyzed the signatures representing these mechanisms, since they can be directly attributed to a known molecular process. All 5 signatures previously associated with MMR and the novel MSI-linked signature N4 exhibit replication strand asymmetry, generally with enrichment of C>T mutations on the leading strand template and C>A and T>C mutations on the lagging strand template (Fig. 4, Fig. 4–figure supplement 1). It has previously been proposed that the correlation of overall mutation rate with replication timing (as shown in Fig 2B) is a direct result of the activity of MMR (Supek & Lehner 2015). In contrast, we observed a more complex relationship. Some MMR signatures in MMR deficient patients do not correlate with replication timing (sig. 15, 21, 26) or do so only in one direction of replication (such as in the leading direction in sig. 20), whereas others show clear timing asymmetry (sig. 6 and N4, Fig. 4– figure supplement 1), indicating that MMR might be only one of several factors influencing mutagenesis in a timing-dependent manner.

**Fig. 4:**
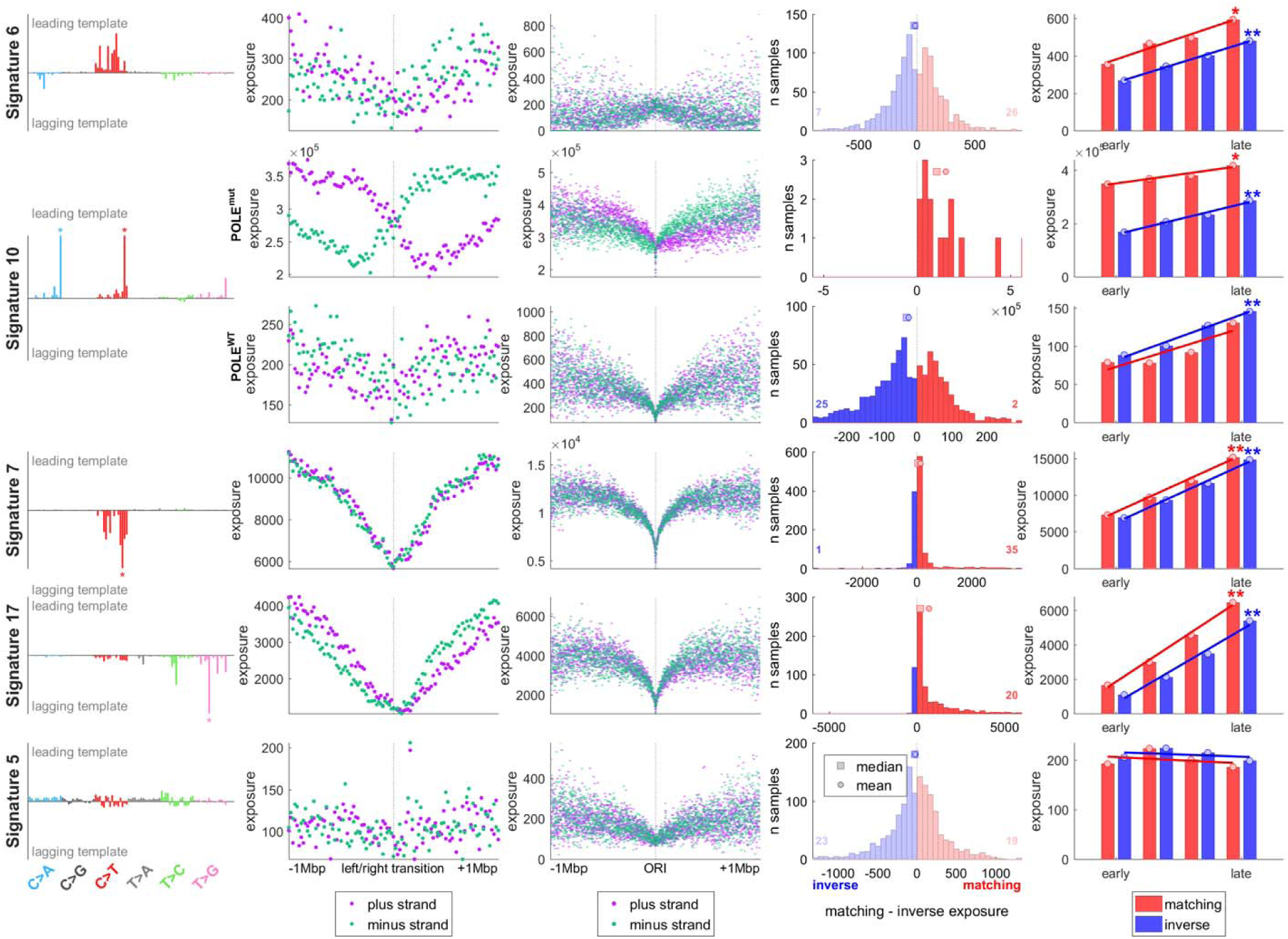
Different mutational signatures exhibit characteristic timing and strand asymmetry profiles. Columns show directional signature (column 1), distribution around timing transition regions (column 2) and around replication origins (column 3), per-patient mutation strand asymmetry (column 4; non-significant asymmetry is shown in light-coloured histogram) and correlation with replication timing (column 5), as described in Fig 3. Row 1: Signature 6, associated with mismatch-repair deficiency. Row 2–3: signature 10, associated with POLE errors, shown for patients with known POLE mutations (row 2), and those without (row 3). Row 4: signature 7, representing UV-induced damage. Row 5: signature 17, characteristic of gastric and esophageal cancers. Row 6: Signature 5, of unknown etiology, is not discernibly affected by replication.

Unexpectedly, two MMR signatures (sig. 6 and N4) showed increased exposures around ORIs (Fig. 4, Fig. 4–figure supplement 1–2, 15). Based on experiments in yeast, it has been suggested that MMR is involved in balancing the differences in fidelity of the leading and lagging polymerases (Lujan et al. 2012), in particular repairing errors made by Pol α (Lujan et al. 2012), which primes the leading strand at ORIs and each lagging strand Okazaki fragment (Stillman 2008) and lacks intrinsic proofreading capabilities (McCulloch & Kunkel 2008). It has been recently shown that error-prone Pol α-synthesized DNA is retained *in vivo*, causing an increase of mutations on the lagging strand (Reijns et al. 2015). Since regions around ORIs have a higher density of Pol α-synthesized DNA (as discussed e.g. in (Waisertreiger et al. 2012)), it is possible that increased exposure to signatures 6 and N4 around ORIs is caused by incomplete repair of Pol α-induced errors. The most common Pol α-induced mismatches normally repaired by MMR are G-dT and C-dT, leading to C>T mutations on the leading and C>A mutations on the lagging strand (Nick McElhinny et al. 2010), matching our observations in the MMR-linked signatures. Notably, we also detected weaker but still significant exposure to MMR signatures in samples with seemingly intact mismatch repair (Fig. 4–figure supplement 2). Replication strand asymmetry in these samples was substantially smaller, but the higher exposure to signatures 6 and N4 around ORIs remained (Fig. 4–figure supplement 15). These findings are compatible with a model in which one of the functions of mismatch repair is to balance the effect of mis-incorporation of nucleotides by Pol α. Signatures 6, N4 and possibly 26 appear to reflect this mechanism, while the other MMR signatures might be a result of unrelated functions of MMR, such as its involvement in balancing errors made by other polymerases, *e.g.* Pol δ.

POLE-M samples were previously reported to be “ultra-hypermutated” with excessive C>A and C>T mutations on the leading strand (Haradhvala et al. 2016; Shinbrot et al. 2014; Shlien et al. 2015). Two mutational signatures (10 and 14) have been associated with Pol ε, the main leading strand polymerase (Stillman 2008; Georgescu et al. 2015). As expected, we observed very strong strand asymmetry for these two signatures in all POLE-M samples, with an increase of C>A, C>T, and T>G mutations on the leading strand (Fig. 4, Fig. 4–figure supplement 3). As with MMR signatures, we also found weak but significant evidence of signature 10 and 14 in samples without Pol ε defects (POLE-WT). Strikingly, however, in these samples the strand asymmetry was in the inverse orientation compared to the POLE-M samples, *i.e.* more C>A, C>T, and T>G mutations on the lagging strand (Fig. 4, Fig. 4–figure supplement 4, 11–14). Conversely, we detected the presence of two signatures of unknown etiology, signatures 18 and 28, in POLE-M samples, but in the inverse orientation compared to POLE-WT samples. In order to validate that this is not an artifact of the signature exposures decomposition, we directly compared the frequencies of the most prominent mutation types for each of the four signatures (sig. 10, 14, 18, and 28) in POLE-MUT and POLE-WT samples on the leading and lagging strands. The inverse strand preference observed in the signatures was also detected for individual mutation types. For example, the frequency of mutations in TCT>A, TCG>T, and TTT>G, the three major components of signature 10, is higher on the lagging strand than on the leading strand in POLE-WT samples, whereas it is higher on the leading strand in POLE-MUT (Fig. 4–figure supplement 11–15). We therefore hypothesize that POLE-linked signatures are originally caused by a process that affects both strands and, under normal circumstances, is slightly enriched on the lagging strand. This could be caused by certain types of DNA lesions which under normal circumstances are less accurately replicated when on the template of the lagging strand (e.g. due to a lower fidelity of Pol δ or Pol α compared to WT Pol ε when replicating these lesions). In POLE-M samples the lack of replication-associated proofreading would then lead to a strong relative increase in these mutations on the leading strand, explaining the flipped orientation of signatures.

### Signatures linked to environmental mutagens

We next focused on signatures that have not previously been reported to be connected to replication, or for which the causal mechanism is unknown. Our data show a link between DNA replication and exogenous mutagens such as UV light (signature 7), tobacco smoke (signature 4) or aristolochic acid (AA) (signature 22) (Helleday et al. 2014). In these signatures, we observed marked correlation with replication timing (Fig. 4, Fig. 4–figure supplement 5–6). Higher mutation frequency late in replication has been observed in mouse embryonic fibroblast (MEFs) treated with AA or Benzo[a]pyrene (B[a]P, a mutagen in tobacco smoke) (Nik-Zainal et al. 2015). This increased mutagenicity might be attributed to different DNA damage tolerance pathways being active during early and late replication. Regions replicated early in S-phase are thought to prefer high-fidelity template switching, whereas regions replicated late are more likely to require translesion synthesis (TLS) which has a higher error rate (Waters & Walker 2006; Lang & Murray 2011; Karras et al. 2013; Gonzalez-Huici et al. 2014; Bi 2015; Branzei & Szakal 2016; D’Souza et al. 2016). This is consistent with the observation in yeast that a disruption of TLS leads to decreased mutation frequency in late-replicating regions and therefore a more even distribution of mutation frequency between early and late-replicating regions (Lang & Murray 2011). In particular, TLS has been observed to increase in activity and mutagenicity later in the cell cycle when replicating DNA damaged by B[a]P (Diamant et al. 2012). Alternatively, differences in chromatin accessibility could be responsible for the decreased mutagenicity in early-replicated regions. Open chromatin is on average replicated earlier and is also more accessible to repair enzymes which could contribute to the decreased mutation frequency in early-replicating regions (Adar et al. 2016).

We also observed weak but significant replication strand asymmetry in the mutagen-induced signatures in the tissues associated with the respective mutagen (Fig. 4–figure supplement 5). Signature 4 has a significant strand asymmetry in lung cancers, similarly as signature 7 in skin cancers. Signature 22 in kidney cancers has small sample size, but shows the same trend. This matches a previously observed lower efficiency of bypass of DNA damage on the lagging strand (Cordeiro-Stone & Nikolaishvili-Feinberg 2002) and strong mutational strand asymmetry in cells lacking Pol η, the main TLS polymerase responsible for the replication of UV-induced photolesions (McGregor et al. 1999). Altogether, our data highlight the importance of replication in converting DNA damage into actual mutations and suggest that bypass of DNA damage occurring on the lagging template results in detectably lower fidelity on this strand.

Signature 17 had the largest median strand asymmetry (p value < 1e^-59^) and also is one of the signatures with the strongest correlations with replication timing (Fig. 2, 4). The mutational process causing this signature is unclear. We noted that the timing asymmetry and exposure distribution around ORIs to signature 17 closely resembled that of signatures 4 and 7, suggesting a possible link to DNA damage. Signature 17 is most prominent in gastric cancers and esophageal adenocarcinoma (EAC), where it appears early during disease development (Murugaesu et al. 2015), and it is also present in Barrett’s esophagus (BE), a precursor to EAC (Ross-Innes et al. 2015). Due to the importance of gastro-esophageal and duodeno-gastric reflux in the development of BE and EAC (Souza 2010; Erichsen et al. 2012; Fein et al. 2006) and the resulting oxidative stress (Kauppi et al. 2016; Rasanen et al. 2007; Jimenez et al. 2005; Dvorak et al. 2007), it has been speculated that oxidative damage could cause the mutation patterns characteristic for Signature 17 (Dulak et al. 2013; Nones et al. 2015). Increased oxidative damage to guanine has been reported in the epithelial cells of dysplastic BE as well as after incubation of BE tissue with a cocktail mimicking bile reflux (Dvorak et al. 2007). Oxidative stress affects not only bases in the DNA, but also the nucleotide pool, such as the oxidation of dGTP to 8-oxo-dGTP. This oxidized dGTP derivative has been shown to induce T>G transversions (Inoue et al. 1998; Satou et al. 2007; Satou et al. 2009) through incorporation by TLS polymerases into DNA opposite A on the template strand (Kamiya 2007). In contrast, oxidation of guanine in the DNA produces 8-oxo-G, which has been shown to result in C>A mutations when paired with adenine during replication (Suzuki & Kamiya 2017). These C>A mutations are normally prevented by DNA glycosylases in the base excision repair pathway, such as MUTYH and OGG1, which repair 8-oxo-G:A pairs to G:C. However, if an 8-oxo-G:A mismatch resulted from incorporation of 8-oxo-dGTP in the de-novo synthesized strand, the “repair” to G:C would actually lead to a T>G mutation (Suzuki & Kamiya 2017). Consequently, depletion of MUTYH lead to an increase of C>A mutations (Suzuki & Kamiya 2017; Rashid et al. 2016) but a decrease of T>G mutations induced by 8-oxo-dGTP (Suzuki et al. 2010). Importantly, the mismatch of 8-oxo-G and A has been shown in yeast to be more efficiently repaired into G:C when 8-oxo-G is on the lagging strand template (Pavlov et al. 2003; Mudrak et al. 2009), resulting in an enrichment of T>G mutations on the lagging strand template if the 8-oxoG:A mismatch originated from incorporation of 8-oxo-dGTP opposite A. Our data show strong lagging-strand bias of T>G mutations and overall higher exposure to signature 17 on the lagging strand, supporting the hypothesis that signature 17 is a by-product of oxidative damage.

## Conclusion

Our findings demonstrate how the relationship between mutational signatures and DNA replication can help to illuminate the mechanisms underlying several currently unexplained mutational processes, as exemplified by Signature 17 in esophageal cancer. Crucially, our computational analysis produces testable hypotheses which we anticipate to be experimentally validated in the future. Our results also add a new perspective to the recent debate regarding the correlation of tissue-specific cell division rates with cancer risk (Tomasetti & Vogelstein 2015). It has been argued that this correlation is primarily attributable to “bad luck” in the form of random errors that are introduced during replication by DNA polymerases. However, the range of mutational signatures observed in cancer samples makes a purely replication-driven etiology of cancer mutations unlikely (Gao et al. 2016; Crossan et al. 2015). Here, we show that most mutational signatures are themselves affected by DNA replication, including signatures linked to environmental mutagens. The presence of mutational signatures one the one hand and a strong relationship between replication and the risk of cancer on the other therefore need not be mutually exclusive. In summary, our results provide evidence that DNA replication interacts with most processes that introduce mutations in the genome, suggesting that differences among DNA polymerases and post-replicative repair enzymes might play a larger part in the accumulation of mutations than previously appreciated.

## MATERIALS AND METHODS

### Somatic mutations

Cancer somatic mutations in 3056 whole-genome sequencing samples (Supplementary Table 1) were obtained from the data portal of The Cancer Genome Atlas (TCGA), the data portal of the International Cancer Genome Consortium (ICGC), and previously published data in peer-review journals (Alexandrov et al. 2013; Bass et al. 2011; Shlien et al. 2015; Dulak et al. 2013; Wang et al. 2014). For the TCGA samples, aligned reads of paired tumor and normal samples were downloaded from the UCSC CGHub website under TCGA access request #10140 and somatic variants were called using Strelka (version 1.0.14) (Saunders et al. 2012) with default parameters.

### Direction of replication

Left- and right-replicating domains were taken from (Haradhvala et al. 2016). Each domain (called territory in the original source code and data) is 20kbp wide and annotated with the direction of replication and with replication timing.

### Excluded regions

The following regions were excluded: regions with low unique mappability of sequencing reads (positions with mean mappability in 100bp sliding windows below 0.99 from UCSC mappability track), gencode protein coding genes, and blacklisted regions defined by Anshul Kundaje (Encode Consortium 2012) (Anshul_Hg19UltraHighSignalArtifactRegions.bed, Duke_Hg19SignalRepeatArtifactRegions.bed, and wgEncodeHg19ConsensusSignalArtifactRegions.bed from http://mitra.stanford.edu/kundaje/akundaje/release/blacklists/hg19-human/).

### Mutation frequency analysis

All variants were classified by the pyrimidine of the mutated Watson-Crick base pair (C or T), strand of this base pair (C or T), and the immediate 5’ and 3’ sequence context into 96 possible mutation types as described by Alexandrov *et al*. (2013). The frequency of trinucleotides on each strand was computed for each replication domain. Then the mutation frequency of each mutation type in each replication domain on the leading (plus=Watson strand in left replicating domains; minus=Crick strand in right replicating domains) and lagging strand (vice versa) was computed for each sample.

### Extraction of mutational signatures

Matlab code (Alexandrov et al. 2013) was used for extraction of strand-specific mutational signatures. The input data were the mutation counts on the leading and lagging strands (summed from all replicating domains together, but without the excluded regions) in each sample. The 192-elements-long mutational signatures (example in Fig. 1B) were extracted in each cancer type separately (for K number of signatures between 2 and 7). The best K with minimal error and maximal stability (minimizing error_K_/max(error) + (1-stability_K_) and with stability of at least 0.8) was selected for each cancer type. Signatures present in only a small number of samples with very low exposures were excluded ((95^th^ percentile of exposures of this signature) / (mean total exposure per samples) < 0.2). The remaining signatures were then normalized by the frequency of trinucleotides in the leading and lagging strand and subsequently multiplied by the frequency of trinucleotides in the genome. This made them comparable with the 30 previously identified whole-genome-based COSMIC signatures (http://cancer.sanger.ac.uk/cosmic/signatures). Signatures extracted in each cancer type and COSMIC signatures were all pooled together (with equal values in the leading and lagging part in the COSMIC signatures) and were clustered using unsupervised hierarchical clustering (with cosine distance and complete linkage). A threshold was selected to identify clusters of similar signatures. Mis-clustering was avoided by manual examination (and whenever necessary re-assignment) of all signatures in all clusters. The resulting 29 signatures (representing the detected clusters) contained 25 previously observed (COSMIC) and 4 new signatures. For the subsequent analysis, the signatures were converted back to 96 values: the 25 previously observed signatures were used in their original form and average of the leading and lagging part were used for the 4 newly identified signatures.

### Annotation of signatures with leading and lagging direction

Each signature was annotated with strand direction: which of the 96 mutation types were higher on the leading strand and which on the lagging strand (Fig. 1C). This was based on the dominant strand direction within the signature’s cluster. Types with unclear direction and small values were assigned according to the predominant direction of other trinucleotides of the same mutation group, such as C>T.

### Calculating strand-specific exposures in individual samples

Exposures to leading and lagging parts of the signatures on the leading and lagging strands in individual samples were quantified using non-negative least squares regression using the Matlab function *e = lsqnonneg(S, m*), where

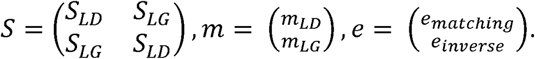

The matrix *S*_*LD*_ has 96 rows and 29 columns and represents the leading parts of the signatures, *i.e.* the elements of the lagging parts contain zeros in this matrix. Similarly, *S*_*LG*_ has the same size, but contains zeros in the leading parts. The vector *m*_*LD*_ of length 96 contains mutations on the leading strand (again normalized by trinucleotides in leading strand/whole genome), and similarly *m*_*LG*_ contains mutations from the lagging strand. Finally, *lsqnonneg* finds a non-negative vector of exposures *e* such that it minimizes a function *|m – S · e|*. A similar approach has been used in (Rosenthal et al. 2016) for finding exposures to a given set of signatures. Our extension includes the strand-specificity of the signatures. The interpretation of the model is that the *matching exposure e*_*matching*_ represents exposure of the leading part of the signature on the leading strand and exposure of the lagging part of the signature on the lagging strand, whereas *e*_*inverse*_ represents the two remaining options. It is important to note that the direction of the mutation is relative to the nucleotide in the base pair chosen as the reference, *i.e.*, mutations of a pyrimidine on the leading strand correspond to mutations of a purine on the lagging strand. In order to minimize the number of spurious signature exposures, the least exposed signature was incrementally removed (in both leading and lagging parts) while the resulting error did not exceed the original error by 0.5%. The resulting reported values in each sample and signature were the difference (or fold change) of *e*_*matching*_ and *e*_*inverse*_. In each signature, the signtest was used to compare matching and inverse exposures across samples with sufficient minimal exposure (at least 10) to the signature. Bonferroni correction was applied to correct for multiple testing.

### Replication origins

The left/right transitions of the replication domains represent regions with on average higher density of replication origins. In order to get better resolution of the replication origins, and to validate the results using an independent estimates of left- and right-replicating domains, genome-wide maps of human replication origins from SNS-seq by (Besnard et al. 2012) were used. Eight fastq files (HeLa, iPS, hESC, IMR; each with two replicates) were downloaded and mapped to hg19 using bowtie2 (version 2.1.0). To control for the inefficient digestion of λ-exo step of SNS-seq, reads from non-replicating genomic DNA (LexoG0) were used as a control (Foulk et al. 2015). Peaks were called using “macs callpeak” with parameters --gsize=hs --bw=200 --qvalue=0.05 --mfold 5 50 and LexoG0 mapped reads as a control. Only peaks covered in at least seven of the eight samples were used. 1000 1kbp bins were generated to the left and right of each origin, as long as they did not reach half the distance to the next origin. We then used these replication direction annotations in the 1kbp bins to calculate strand-specific exposures in individual samples as above and ascertained that both approaches lead to qualitatively very similar mutational strand asymmetries in individual signatures (Fig. 2–figure supplement 4, 7).

### Quantification of exposures with respect to replication timing, left/right transitions, and replication origins

Replication domains were divided into four quartiles by their average replication timing. The entire exposure quantification was computed separately in each quartile, or bin around left/right transition or bin around replication origin. In replication timing plots, a linear regression model (function `fitlm` in MatLab) was fitted to the mean exposure in each quartile (separately for matching and inverse exposures) and the significance of the linear coefficient was tested using F-test for the hypothesis that the regression coefficient is zero (function `coefTest` in MatLab).

## Footnotes

1 http://cancer.sanger.ac.uk/cosmic/signatures

## ACKNOWLEDGMENTS

We thank Dr. Mary Muers for comments on the manuscript. S.K. and B.S.-B. are funded by Ludwig Cancer Research. S.K. received funding from BBSRC grant BB/M001873/1. M.T. and J.T. are funded by EPSRC (EP/F500394/1) and Bakala Foundation. Author contributions: B.S.-B. and M.T. designed the study; M.T. performed the analysis with contributions from J.T.; B.S.-B. and M.T. wrote the manuscript with contributions from S.K. and J.T.

## REFERENCES

Adar, S. et al., 2016. Genome-wide kinetics of DNA excision repair in relation to chromatin state and mutagenesis. Proceedings of the National Academy of Sciences of the United States of America, p.201603388. Available at: http://www.pnas.org/lookup/doi/10.1073/pnas.1603388113%5Cnhttp://www.ncbi.nlm.nih.gov/pubmed/27036006%5Cn http://www.pubmedcentral.nih.gov/articlerender.fcgi?artid=PMC4839430.

Alexandrov, L.B. et al., 2013. Signatures of mutational processes in human cancer. Nature, 500(7463), pp.415–21. Available at: http://www.pubmedcentral.nih.gov/articlerender.fcgi?artid=3776390&tool=pmcentrez&rendertype=abstract [Accessed March 19, 2014].

Bass, A.J. et al., 2011. Genomic sequencing of colorectal adenocarcinomas identifies a recurrent VTI1ATCF7L2 fusion. Nature genetics, 43(10), pp.964–8. Available at: http://www.pubmedcentral.nih.gov/articlerender.fcgi?artid=3802528&tool=pmcentrez&rendertype=abstract.

Besnard, E. et al., 2012. Unraveling cell type-specific and reprogrammable human replication origin signatures associated with G-quadruplex consensus motifs. Nature structural & molecular biology, 19(8), pp.837–844.

Bi, X., 2015. Mechanism of DNA damage tolerance. World Journal of Biological Chemistry, 6(3), p.48. Available at: http://www.wjgnet.com/1949-8454/full/v6/i3/48.htm.

Branzei, D. & Szakal, B., 2016. DNA damage tolerance by recombination: Molecular pathways and DNA structures. DNA Repair, 44, pp.68–75. Available at: http://dx.doi.org/10.1016/j.dnarep.2016.05.008.

Cordeiro-Stone, M. & Nikolaishvili-Feinberg, N., 2002. Asymmetry of DNA replication and translesion synthesis of UV-induced thymine dimers. Mutation research, 510(1-2), pp.91–106.

Crossan, G.P., Garaycoechea, J.I. & Patel, K.J., 2015. Do mutational dynamics in stem cells explain the origin of common cancers? Cell Stem Cell, 16(2), pp.111–112. Available at: http://dx.doi.org/10.1016/j.stem.2015.01.009.

D’Souza, S., Yamanaka, K. & Walker, G.C., 2016. Non mutagenic and mutagenic DNA damage tolerance. Cell Cycle, 15(3), pp.314–315. Available at: http://dx.doi.org/10.1080/15384101.2015.1132909.

Diamant, N. et al., 2012. DNA damage bypass operates in the S and G2 phases of the cell cycle and exhibits differential mutagenicity. Nucleic Acids Research, 40(1), pp.170–180.

Dulak, A.M. et al., 2013. Exome and whole-genome sequencing of esophageal adenocarcinoma identifies recurrent driver events and mutational complexity. Nature genetics, 45(5), pp.478–86. Available at: http://www.pubmedcentral.nih.gov/articlerender.fcgi?artid=3678719&tool=pmcentrez&rendertype=abstract.

Dvorak, K. et al., 2007. Bile acids in combination with low pH induce oxidative stress and oxidative DNA damage: relevance to the pathogenesis of Barrett’s oesophagus. Gut, 56, pp.763–771.

Encode Consortium, 2012. An integrated encyclopedia of DNA elements in the human genome. Nature, 489(7414), pp.57–74. Available at: http://www.nature.com/nature/journal/v489/n7414/full/nature11247.html%5Cnhttp://www.nature.com/nature/journal/v489/n7414/pdf/nature11247.pdf.

Erichsen, R. et al., 2012. Erosive Reflux Disease Increases Risk for Esophageal Adenocarcinoma, Compared With Nonerosive Reflux. Clinical Gastroenterology and Hepatology, 10(5), p.475–480.e1. Available at: http://dx.doi.org/10.1016/j.cgh.2011.12.038.

Fein, M., Maroske, J. & Fuchs, K.H., 2006. Importance of duodenogastric reflux in gastro-oesophageal reflux disease. British Journal of Surgery, 93(12), pp.1475–1482.

Foulk, M.S. et al., 2015. Characterizing and controlling intrinsic biases of lambda exonuclease in nascent strand sequencing reveals phasing between nucleosomes and G-quadruplex motifs around a subset of human replication origins. Genome Research, 25, pp.725–735.

Fragkos, M. et al., 2015. DNA replication origin activation in space and time. Nature reviews. Molecular cell biology, 16(6), pp.360–74. Available at: http://dx.doi.org/10.1038/nrm4002%5Cnhttp://www.ncbi.nlm.nih.gov/pubmed/25999062.

Gao, Z. et al., 2016. Interpreting the Dependence of Mutation Rates on Age and Time N. H. Barton, ed. PLoS Biology, 14(1), p.e1002355. Available at: http://dx.plos.org/10.1371/journal.pbio.1002355 [Accessed March 10, 2017].

Georgescu, R.E. et al., 2015. Reconstitution of a eukaryotic replisome reveals suppression mechanisms that define leading/lagging strand operation. eLife, 2015(4), pp.1–20.

Gonzalez-Huici, V. et al., 2014. DNA bending facilitates the error-free DNA damage tolerance pathway and upholds genome integrity. The EMBO Journal, 33(4), pp.327–340.

Green, A.M. et al., 2016. APOBEC3A damages the cellular genome during DNA replication. Cell Cycle, 15(7), pp.998–1008. Available at: http://dx.doi.org/10.1080/15384101.2016.1152426.

Haradhvala, N.J. et al., 2016. Mutational Strand Asymmetries in Cancer Genomes Reveal Mechanisms of DNA Damage and Repair. Cell, 164(3), pp.538–549. Available at: http://linkinghub.elsevier.com/retrieve/pii/S0092867415017146.

Helleday, T., Eshtad, S. & Nik-Zainal, S., 2014. Mechanisms underlying mutational signatures in human cancers. Nature reviews. Genetics, 15(9), pp.585–598. Available at: http://www.ncbi.nlm.nih.gov/pubmed/24981601.

Hoopes, J.I. et al., 2016. APOBEC3A and APOBEC3B Preferentially Deaminate the Lagging Strand Template during DNA Replication. Cell Reports, pp.1–10. Available at: http://linkinghub.elsevier.com/retrieve/pii/S2211124716000425.

Inoue, M. et al., 1998. Induction of chromosomal gene mutations in Escherichia coli by direct incorporation of oxidatively damaged nucleotides: New evaluation method for mutagenesis by damaged dna precursors in vivo. Journal of Biological Chemistry, 273(18), pp.11069–11074.

Jimenez, P. et al., 2005. Free radicals and antioxidant systems in reflux esophagitis and Barrett’s esophagus. World journal of gastroenterology: WJG, 11(18), pp.2697–2703. Available at: http://www.ncbi.nlm.nih.gov/pubmed/15884106.

Kamiya, H., 2007. Mutations Induced by Oxidized DNA Precursors and Their Prevention by Nucleotide Pool Sanitization Enzymes. Genes and Environment, 29(4), pp.133–140.

Karras, G.I. et al., 2013. Article Noncanonical Role of the 9-1-1 Clamp in the Error-Free DNA Damage Tolerance Pathway. Molecular Cell, 49(3), pp.536–546. Available at: http://dx.doi.org/10.1016/j.molcel.2012.11.016.

Kauppi, J. et al., 2016. Increased Oxidative Stress in the Proximal Stomach of Patients with Barrett’s Esophagus and Adenocarcinoma of the Esophagus and Esophagogastric Junction. Translational oncology, 9(4), pp.336–339. Available at: http://www.ncbi.nlm.nih.gov/pubmed/27567957.

Lang, G.I. & Murray, A.W., 2011. Mutation rates across budding yeast chromosome VI Are correlated with replication timing. Genome Biology and Evolution, 3(1), pp.799–811.

Lawrence, M.S. et al., 2013. Mutational heterogeneity in cancer and the search for new cancer-associated genes. Nature, 499(7457), pp.214–8. Available at: http://dx.doi.org/10.1038/nature12213 [Accessed July 11, 2014].

Lujan, S.A. et al., 2012. Mismatch Repair Balances Leading and Lagging Strand DNA Replication Fidelity. PLoS Genetics, 8(10), p.e1003016.

Lujan, S.A., Williams, J.S. & Kunkel, T.A., 2016. DNA Polymerases Divide the Labor of Genome Replication. Trends in Cell Biology, 26(9), pp.640–654. Available at: http://dx.doi.org/10.1016/j.tcb.2016.04.012.

McCulloch, S.D. & Kunkel, T.A., 2008. The fidelity of DNA synthesis by eukaryotic replicative and translesion synthesis polymerases. Cell Research, 18, pp.148–161.

McGregor, W.G. et al., 1999. Abnormal, Error-Prone Bypass of Photoproducts by Xeroderma Pigmentosum Variant Cell Extracts Results in Extreme Strand Bias for the Kinds of Mutations Induced by UV Light. Molecular and Cellular Biology, 19(1), pp.147–154. Available at: http://mcb.asm.org/content/19/1/147.abstract.

Morganella, S. et al., 2016. The topography of mutational processes in breast cancer genomes. Nature Communications, 7(may 2016), p.11383. Available at: http://www.nature.com/doifinder/10.1038/ncomms11383.

Mudrak, S. V, Welz-Voegele, C. & Jinks-Robertson, S., 2009. The polymerase eta translesion synthesis DNA polymerase acts independently of the mismatch repair system to limit mutagenesis caused by 7,8-dihydro-8-oxoguanine in yeast. Molecular and cellular biology, 29(19), pp.5316–26. Available at: http://www.scopus.com/inward/record.url?eid=2-s2.0-70349329456&partnerID=tZOtx3y1.

Murugaesu, N. et al., 2015. Tracking the genomic evolution of esophageal adenocarcinoma through neoadjuvant chemotherapy. Cancer Discovery, 5(8), pp.821–832.

Nick McElhinny, S. a, Kissling, G.E. & Kunkel, T. a, 2010. Differential correction of lagging-strand replication errors made by DNA polymerases a and d. Proceedings of the National Academy of Sciences of the United States of America, 107(49), pp.21070–21075.

Nik-Zainal, S. et al., 2015. The genome as a record of environmental exposure. Mutagenesis, 30(October), pp.763–770.

Nones, K. et al., 2015. Genomic catastrophes frequently arise in esophageal adenocarcinoma and drive tumorigenesis. Nature Communications, 5, pp.1–9. Available at: http://dx.doi.org/10.1038/ncomms6224%5Cnpapers2://publication/doi/10.1038/ncomms6224.

Pavlov, Y.I., Mian, I.M. & Kunkel, T.A., 2003. Evidence for Preferential Mismatch Repair of Lagging Strand DNA Replication Errors in Yeast. Current Biology, 13, pp.744–748.

Rasanen, J. V. et al., 2007. The expression of 8-hydroxydeoxyguanosine in oesophageal tissues and tumours. European Journal of Surgical Oncology, 33(10), pp.1164–1168.

Rashid, M. et al., 2016. Adenoma development in familial adenomatous polyposis and MUTYH-associated polyposis: Somatic landscape and driver genes. Journal of Pathology, 238(1), pp.98–108.

Reijns, M.A.M. et al., 2015. Lagging-strand replication shapes the mutational landscape of the genome. Nature, 518(7540), pp.502–506. Available at: http://www.nature.com/nature/journal/v518/n7540/full/nature14183.html%5Cnhttp://www.nature.com/nature/journal/v518/n7540/pdf/nature14183.pdf.

Rosenthal, R. et al., 2016. deconstructSigs: delineating mutational processes in single tumors distinguishes DNA repair deficiencies and patterns of carcinoma evolution. Genome Biology, 17(1), p.31. Available at: http://genomebiology.biomedcentral.com/articles/10.1186/s13059-016-0893-4.

Ross-Innes, C.S. et al., 2015. Whole-genome sequencing provides new insights into the clonal architecture of Barrett’s esophagus and esophageal adenocarcinoma. Nature Genetics, 47(July), pp.1–11. Available at: http://www.nature.com/doifinder/10.1038/ng.3357.

Satou, K. et al., 2009. Involvement of specialized DNA polymerases in mutagenesis by 8-hydroxy-dGTP in human cells. DNA Repair, 8(5), pp.637–642.

Satou, K. et al., 2007. Mutagenic effects of 8-hydroxy-dGTP in live mammalian cells. Free Radical Biology and Medicine, 42(10), pp.1552–1560. Available at: http://dx.doi.org/10.1016/j.freeradbiomed.2007.02.024.

Saunders, C.T. et al., 2012. Strelka: Accurate somatic small-variant calling from sequenced tumor-normal sample pairs. Bioinformatics, 28(14), pp.1811–1817.

Secrier, M. et al., 2016. Mutational signatures in esophageal adenocarcinoma define etiologically distinct subgroups with therapeutic relevance. Nature Genetics, 2016(September), pp.1131–1141. Available at: http://www.nature.com/doifinder/10.1038/ng.3659.

Seplyarskiy, V.B. et al., 2016. APOBEC-induced mutations in human cancers are strongly enriched on the lagging DNA strand during replication. Genome Research, 26(2), pp.174–182.

Shinbrot, E. et al., 2014. Exonuclease mutations in DNA Polymerase epsilon reveal replication strand specific mutation patterns and human origins of replication. Genome Research, pp.1740–1750. Available at: http://www.ncbi.nlm.nih.gov/pubmed/25228659.

Shlien, A. et al., 2015. Combined hereditary and somatic mutations of replication error repair genes result in rapid onset of ultra-hypermutated cancers. Nature Genetics, 47(3), pp.257–262. Available at: http://www.nature.com/doifinder/10.1038/ng.3202.

Souza, R.F., 2010. The role of acid and bile reflux in oesophagitis and Barrett’s metaplasia. Biochemical Society transactions, 38(2), pp.348–52. Available at: http://www.pubmedcentral.nih.gov/articlerender.fcgi?artid=3072824&tool=pmcentrez&rendertype=abstract.

Stamatoyannopoulos, J. a et al., 2009. Human mutation rate associated with DNA replication timing. Nature genetics, 41(4), pp.393–395.

Stenzinger, A. et al., 2014. Mutations in POLE and survival of colorectal cancer patients--link to disease stage and treatment. Cancer medicine, 3(6), pp.1527–1538.

Stillman, B., 2008. DNA Polymerases at the Replication Fork in Eukaryotes. Molecular Cell, 30(3), pp.259–260.

Supek, F. & Lehner, B., 2015. Differential DNA mismatch repair underlies mutation rate variation across the human genome. Nature, 521(7550), pp.81–84.

Suzuki, T., Harashima, H. & Kamiya, H., 2010. Effects of base excision repair proteins on mutagenesis by 8-oxo-7,8-dihydroguanine (8-hydroxyguanine) paired with cytosine and adenine. DNA Repair, 9(5), pp.542–550. Available at: http://dx.doi.org/10.1016/j.dnarep.2010.02.004.

Suzuki, T. & Kamiya, H., 2017. Mutations induced by 8-hydroxyguanine (8-oxo-7,8-dihydroguanine), a representative oxidized base, in mammalian cells. Genes and Environment, 12, pp.4–9. Available at: http://dx.doi.org/10.1186/s41021-016-0051-y.

Tomasetti, C. & Vogelstein, B., 2015. Variation in cancer risk among tissues can be explained by the number of stem cell divisions. Science (New York, N.Y.), 347(6217), pp.78–81. Available at: http://www.ncbi.nlm.nih.gov/pubmed/25554788.

Waisertreiger, I.S.-R. et al., 2012. Modulation of Mutagenesis in Eukaryotes by DNA Replication Fork Dynamics and Quality of Nucleotide Pools. Environmental and molecular mutagenesis, 53(9), pp.699–724. Available at: http://www.ncbi.nlm.nih.gov/pubmed/23055184 [Accessed May 2, 2017].

Wang, K. et al., 2014. Whole-genome sequencing and comprehensive molecular profiling identify new driver mutations in gastric cancer. Nature genetics, 46(6), pp.573–82. Available at: http://www.ncbi.nlm.nih.gov/pubmed/24816253.

Waters, L.S. & Walker, G.C., 2006. The critical mutagenic translesion DNA polymerase Rev1 is highly expressed during G(2)/M phase rather than S phase. Proceedings of the National Academy of Sciences of the United States of America, 103(24), pp.8971–8976.

Zhao, H. et al., 2014. Mismatch repair deficiency endows tumors with a unique mutation signature and sensitivity to DNA double-strand breaks. eLife, 3, p.e02725.

